# Chronic hypertension disrupts the blood-brain barrier in the ventral hippocampus in females but not males in a rat model of vascular cognitive impairment

**DOI:** 10.64898/2026.10.31.763337

**Authors:** Jared Luchetta, Gianna M. DeMarco, Anduela Thaqi, Gabrielle A. Bernstein, Swapna E. Chavara, Emily D. Schellinger, Dinlin Chen, Abbie C. Chapman

**Affiliations:** Department of Psychology, Binghamton University – State University of New York, Binghamton, NY, USA 13902

**Keywords:** anxiety-related behavior, blood-brain barrier, chronic hypertension, sex differences, vascular cognitive impairment, ventral hippocampus

## Abstract

**Background:** Chronic hypertension is a leading cause of cerebrovascular disease, ischemic stroke, and cognitive impairment. In addition, anxiety and hypertension often co-exist, however, little is known about the relationship between the two. Understanding how chronic hypertension impacts limbic brain regions such as the ventral hippocampus may provide insight into how cerebrovascular contributes to altered emotional behavior.

**Methods:** An open field task was used to quantify locomotion, exploratory behavior, thigmotaxis, rearing and risk assessment behaviors in adult (7-month-old) male and female normotensive Wistar rats and spontaneously hypertensive rats (SHR; n=8/group). Blood-brain barrier (BBB) permeability was assessed by measuring extravasation of sodium fluorescein, 3 kDa Texas red dextran and 10 kDa cascade blue into the ventral hippocampus. Enzyme-linked immunosorbent assays were used to measure markers of vascular signaling, inflammation, and central nervous system-specific proteins indicative of neuroaxonal and glial injury in serum from separate groups of male and female SHR (n=6/group). Main effects and interactions between two independent variables (sex and hypertension) were determined using two-way analysis of variance (ANOVA) with Tukey’s post-hoc tests. Circulating factors were compared using a Welch’s t-test. Relationships between BBB permeability and anxiety-like behavior were determined using Pearson’s correlations.

**Results:** Chronic hypertension increased circulating inflammatory factors and BBB permeability in female but not male rats. BBB permeability did not differ by sex in male and female normotensive rats. There was an anxiolytic effect of chronic hypertension on exploratory-related anxiety-like behavior, including less thigmotaxis, lower latency to cross through the center of the arena, and increased unsupported rearing. Risk assessment behavior, quantified as stretch-attend posture, was increased in female SHR. BBB permeability in the ventral hippocampus was correlated with anxiety-like behavior in females, but not in males, suggesting sex-dependent mechanisms of hypertension-induced changes in emotional behavior.

**Conclusions:** These findings implicate a role for chronic hypertension-induced BBB disruption in neuropsychiatric disorders in the female sex and advances our understanding of cerebrovascular vulnerability in deep limbic brain regions.

## Introduction

Chronic hypertension is a leading cause of cerebrovascular disease, ischemic stroke, and vascular contributions to cognitive impairment and dementia.^1^ Persistently elevated blood pressure (> 130 mmHg systolic / 80 mmHg diastolic) is considered a “silent killer” because it is often asymptomatic and remain undiagnosed.^2, 3^ Even when hypertension is diagnosed, blood pressure often remains inadequately controlled, allowing chronic vascular stress to persist.^4, 5^ Although hypertension is an established risk factor for dementia,^6^ its potential contribution to neuropsychiatric outcomes has received comparatively less attention. This gap is particularly important for anxiety disorders, which represent a major global health burden and are regulated by brain regions affected by hypertension. The Global Burden of Disease Mental Health Disorders Collaborators reported 470 million cases of anxiety disorders worldwide in 2023, with anxiety representing a major reason for loss of healthy life years globally.^7^ If hypertension-induced cerebrovascular damage contributes to anxiety-related outcomes, then reversing this damage could potentially alter the severity or persistence of anxiety disorders. However, the extent to which hypertension contributes to anxiety, and through what mechanisms, remains poorly understood.

Chronic hypertension is known to damage several critical aspects of cerebrovascular structure and function, including the blood-brain barrier (BBB). The BBB is a highly protective interface between blood constituents circulating systemically and the delicate microenvironment of the brain. The innermost layer of the BBB is the vascular endothelium. In the brain, this layer is unique and only allows selective passage of solutes, proteins, cells, etc. from blood to brain (and to a lesser extent vice versa).^8^ The BBB has this capability because of specialized high electrical-resistance tight junctions that anchor endothelial cells of the BBB to one another; in conjunction with the lack of fenestrations, they limit para- and transcellular permeability.^9^ Disruption of the BBB can cause neuroinflammation, neuronal dysfunction, and impaired cognition. BBB breakdown is considered an early biomarker of human cognitive decline.^1, 10, 11^ Females in general have lower BBB permeability than males, including in the hippocampus, which is attributed to protective actions of estrogens and higher expression of tight junction proteins.^12, 13^ However, age-related increases in BBB permeability occur more rapidly in males than females, suggesting conditions such as chronic hypertension may damage the BBB in a sex-specific manner.^14^

Anxiety serves to detect and avoid threats, and either excessive or diminished responses can be detrimental to an individual’s well-being. Previous studies have reported that chronic hypertension has a general anxiolytic effect, specifically in a commonly used rat model of vascular cognitive impairment and dementia, the spontaneously hypertensive rat (SHR).^15, 16^ However, it remains unclear whether chronic hypertension produces sex-specific BBB disruption within the ventral hippocampus, a region centrally involved in emotional processing and anxiety-related behavior, or whether BBB permeability is associated with hypertension-induced changes in anxiety-like behavior.^17^ In the current study, we investigated the effects of sex and chronic hypertension on BBB permeability in the ventral hippocampus *in vivo* and its relationship with anxiety-like behavior in adult male and female normotensive and chronically hypertensive rats. We hypothesized that BBB permeability would be higher in the ventral hippocampus during hypertension in males than in females, and that this would be correlated with anxiety-like behavior.

## Methods

### Animals

Male and female Wistar rats and SHR were purchased from Charles River at 3 months of age (N=24; n=8/group) and housed in the Binghamton University animal facility, an Association for Assessment and Accreditation of Laboratory Animal Care International (AAALAC) accredited facility. Animals were maintained on a 12-h light/dark cycle with access to food and water ad libitum. All procedures were approved by the Binghamton University Institutional Animal Care and Use Committee (protocol #s 898-23 and 956-25) and conducted in accordance with the NIH Guide for the Care and Use of Laboratory Animals. Euthanasia occurred via exsanguination followed by rapid decapitation under isoflurane anesthesia (3% in oxygen).

### BBB permeability in vivo

To understand how sex and chronic hypertension impact the vasculature of limbic brain regions, BBB permeability to three different-sized tracers: 0.3 kDa sodium fluorescein (NaFl, Stokes-Einstein radius ∼ 0.45nm; 5mg/mL), 3 kDa Texas red (Stokes-Einstein radius ∼ 1.3nm; 1mg/mL) and 10 kDa cascade blue (Stokes-Einstein radius ∼ 2.35nm; 1mg/mL) was measured *in vivo* in the ventral hippocampus.^18, 19^ Under isoflurane anesthesia, the tracer cocktail was injected via tail vein (1.5mL at 5 mL/min) and allowed to circulate for 5 minutes. The circulation was then flushed with cold 0.1M PBS (∼ 80mL, 10mL/min) via transcardiac perfusion. All tracer stocks and cocktails were protected from light. Under red light, brains were removed and the ventral hippocampus dissected, weighed and homogenized using a bead homogenizer for two minutes (2 m/s in 800uL of 0.1M PBS). Tissue homogenates were deproteinated with 800µL of 50% TCA (trichloroacetic acid) and vortexed for one minute. Samples were centrifuged (10 minutes at 4500 rpm and 4°C), supernatants collected and centrifuged again. Samples were plated in triplicate in black 96-well plates. TCA/PBS wells, consisting of an equal mixture of 50% TCA and 0.1M PBS were used as blank controls. Fluorescence was quantified as relative fluorescence unit (RFU) using a plate reader (Tecan Infinite 200 Pro). Background fluorescence from TCA/PBS wells was subtracted and values normalized to tissue weight. Triplicate values were averaged per sample. Composite scores were generated by summating permeability normalized to weight for all three tracers to represent a total BBB Score for each animal. Pilot studies were conducted to determine tracer concentrations and circulation time needed to detect tracer passage into the brain and kidney was used as a positive control, as previously described.^18^

### Behavioral assessments

At 6 months old, an open field arena task (5 min) was used to analyze locomotion, exploratory behavior and anxiety-like behavior prior to BBB permeability measurements. Automated tracking software AnyMaze (Stoelting Co., Wood Dale, IL, USA) was used to quantify speed, distance traveled, latency to cross through the center of the arena, number of center crossings, and thigmotaxis, e.g., the time spent in perimeter/middle/center areas of the arena. The number of and time spent in 1) supported rearing – rearing onto the hindlimbs with one or both forepaws on the arena wall and considered general exploratory behavior, 2) unsupported rearing – rearing onto the hindlimbs without contact with the arena wall, which is a behavior highly sensitive to anxiety,^20^ and 3) stretch-attend posture – body elongation towards the center with paws planted in the perimeter of the arena as risk assessment during an approach-avoidant conflict that is heavily dependent upon the ventral hippocampus^21^ were quantified by two independent investigators that were blinded to group. The open field arena (MazeEngineers, Skokie, IL, USA) was cleaned with 70% ethanol between animals. All rats acclimated to the behavior room for one hour before behavior began and all open field tasks were run at the same time of day.

### Measurement of circulating factors

In separate groups of age-matched male and female SHR (n=6/group), blood was collected via cardiac puncture, centrifuged for 10 minutes at room temperature, and serum aliquoted and stored at −80°C until use. Circulating factors that damage the BBB were measured via commercially available enzyme-linked immunosorbent assays (ELISAs). More specifically, ELISAs were used to quantify serum levels of VEGF (vascular endothelial growth factor, Product RRV00; samples undiluted), ICAM-1 (intercellular adhesion molecule 1, Product RIC100; sample run with 50-fold dilution) and MMP-9 (matrix metalloproteinase-9, product RMP900; samples undiluted) obtained from R&D Systems (Minneapolis, MN, USA) and IL-17A (interleukin-17A, Product ab119536, samples run with 2-fold dilution) purchased from Abcam (Waltham, MA, USA). To assess astrocyte activation and neuroaxonal injury alongside BBB permeability, we measured serum GFAP (glial fibrillary acidic protein, Product MBS2505953, samples undiluted) purchased from MyBiosource, Inc. (San Diego, CA, USA) and NF-L (neurofilament light chain, Product NBP2-81185, samples undiluted) from Novus Biological (Centennial, CO, USA). All samples were randomized across plates and run in duplicate according to manufacturer instructions.

### Drugs and solutions

All tracers were diluted in sterile Lactated Ringer’s solution (Baxter, Deerfield, IL, USA) and cocktails prepared fresh daily. Stock tracer solutions were protected from light and stored according to manufacturer’s recommendations. 0.1 M PBS was prepared weekly and stored at 4 °C. A 50% TCA (Sigma-Aldrich, St. Louis, MO, USA) stock solution was prepared by dissolving TCA in Milli-Q water, aliquoted, and stored at 4 °C. Working solutions were made fresh as needed for protein precipitation during tissue processing. All buffer solutions were prepared weekly and stored at 4 °C until use.

### Statistical analysis

Sample size was based on a statistical power calculation to determine the number of animals needed using a two-sided 95% confidence interval for a single mean and 1 – β of 0.80 based on our prior studies using similar methodology.^19, 22, 23^ Through pilot studies, we determined that n=6/group was sufficient to detect differences in passage of fluorescent tracers into the brain and locomotion/anxiety-like behavior. Based on an 80% success rate in pilot permeability studies, we added two rats to each group, totaling n=8/group. Main effects and the interaction between sex and hypertension/housing condition were determined using a two-way ANOVA with a Tukey post-hoc test to correct for multiple comparisons. Linear regression analyses were generated to understand the relationships between BBB permeability and anxiety-like behavior. Welch’s t-test was used to compare serum levels of circulating factors between male and female SHR. The normality assumption of residuals was assessed via visual inspection of QQ plots showing residuals closely followed the reference line and supported by non-significant Shapiro-Wilk tests. Results are reported as mean ± SEM, with p < 0.05 considered significant. All analyses were performed in GraphPad Prism 10.3. Tissue processing and BBB permeability analyses were completed by investigators who were blinded to group.

## Results

Chronic hypertension is associated with increased anxiety in women compared to men, however, how it impacts cerebrovascular structure and function within limbic brain regions remain unclear. In the current study, we measured BBB permeability *in vivo* to three sized tracers in the ventral hippocampus to understand how chronic hypertension disrupted barrier function and whether this differed by sex. There were main effects of sex and hypertension, and a significant interaction between the two on body weight (F_1,28_ = 886.3; p<0.001; F_1,28_ = 244.1; p<0.001; F_1,28_ = 25.33; p<0.001 by two-way ANOVA, respectively). Males weighed more than females, and chronic hypertension decreased body weights in both sexes (male Wistar: 532 ± 8 g, male SHR: 389 ± 8 g, female Wistar: 291 ± 6 g, female SHR: 246 ± 3 g by two-way ANOVA with post-hoc Tukey’s test).

Figure 1 shows BBB permeability to ∼ 400 Da sodium fluorescein (NaFl; Figure 1A), 3 kDa Texas reg (TxR; Figure 1B), 10 kDa cascade blue (CasB; Figure 1C) and composite scores of total BBB permeability (Figure 1D). Extravasation of tracers through the BBB decreased as the size of tracer increased. There were main effects of sex and hypertension, but no significant interaction on NaFl permeability (F_1,28_ = 14.68, p=0.0007; F_1,28_ = 8.758, p=0.0062; F_1,28_ = 3.431, p=0.0746 by two-way ANOVA, respectively). BBB permeability was higher in female SHR compared to female Wistars and male SHR (Figure 1A; p<0.01, p<0.05 by two-way ANOVA with a post-hoc Tukey test). There was a main effect of hypertension, but not of sex, and no interaction between the two on permeability to 3 kDa TxR (F_1,25_ = 6.858, p=0.0148; F_1,25_ = 0.2724, p=0.6063; F_1,25_ = 2.455, p=0.1297 by two-way ANOVA, respectively). There was no effect of chronic hypertension on permeability to 3 kDa TxR in the ventral hippocampus in males, however, hypertension significantly increased permeability in females (Figure 1B; p<0.05 by two-way ANOVA with post-hoc Tukey test). There were main effects of sex and hypertension, but no interaction between sex and hypertension on permeability to 10 kDa CasB (F_1,28_ = 20.48, p=0.0001; F_1,28_ = 4.283, p=0.0478; F_1,28_ = 3.073, p=0.0906 by two-way ANOVA, respectively). Permeability was higher in female SHR compared to normotensive Wistar rats and male SHR (p=0.053, p<0.01 by two-way ANOVA with post-hoc Tukey test; Figure 1C). There were main effects of sex and hypertension and a significant sex-by-hypertension interaction in total BBB permeability, as shown by composite BBB scores (F_1,27_ = 16.23, p=0.0004; F_1,27_ = 12.41, p=0.0015; F_1,27_ = 8.730, p=0.0064 by two-way ANOVA, respectively; Figure 1D). BBB permeability in the ventral hippocampus was unaffected by chronic hypertension in males, however, was higher in female SHR compared to male SHR and normotensive female rats (p<0.01 by two-way ANOVA with post-hoc Tukey test; Figure 1D). One female Wistar rat was excluded from 3 kDa TxR measurements and, therefore, from the total BBB score due to being a statistical outlier (> 2SD from the mean). These data suggest that BBB permeability in the ventral hippocampus to solutes ≤ 10 kDa in size was similar between normotensive males and females. They further suggest that chronic hypertension caused disruption of the BBB in the ventral hippocampus in females but had little-to-no effect in males.

**Figure 1.**
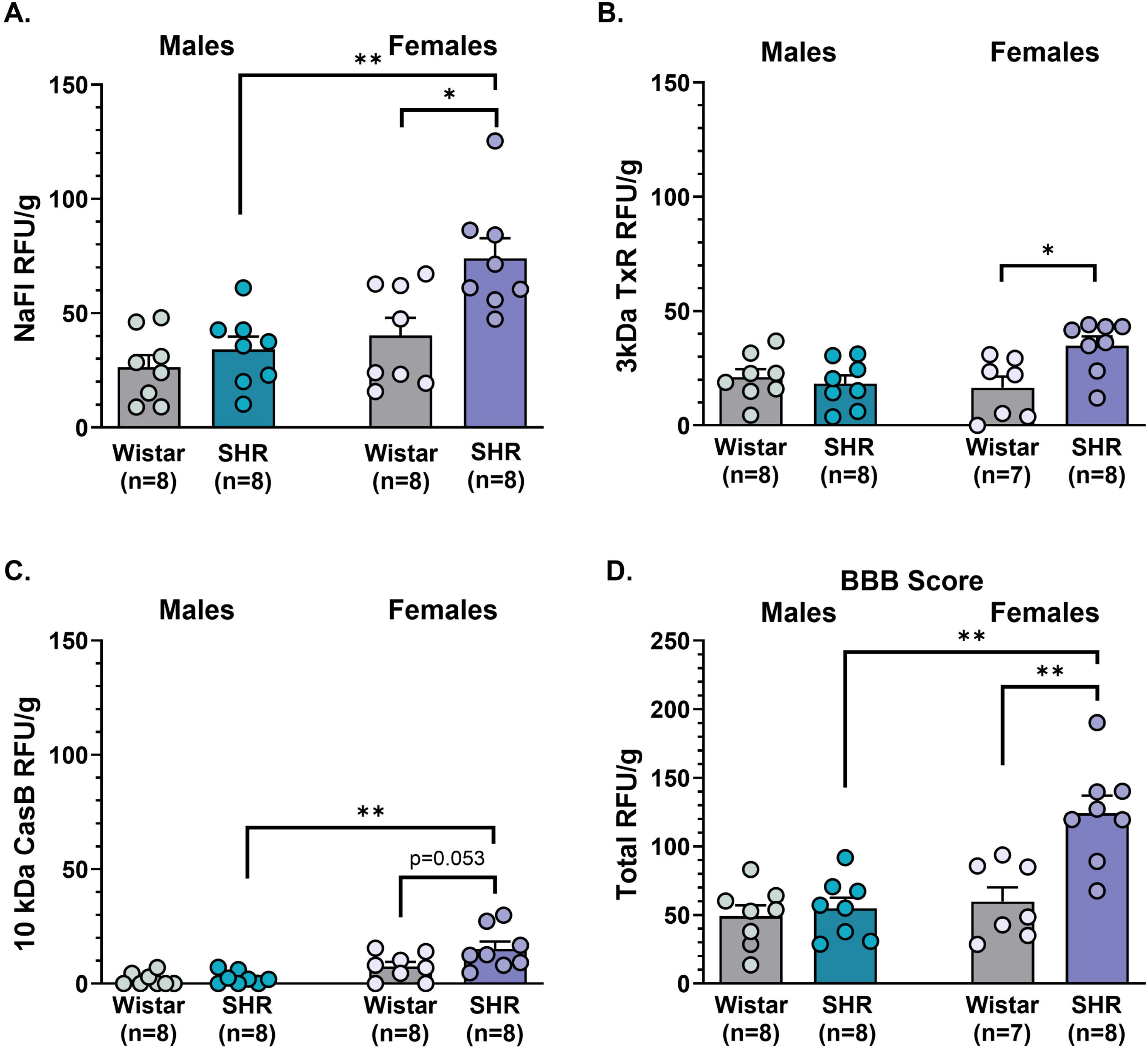
Blood-brain barrier (BBB) permeability in ventral hippocampus during chronic hypertension. BBB permeability to **A.** small solute sodium fluorescein (NaFl), **B**. 3 kDa Texas red (TxR), **C.** 10 kDa cascade blue (CasB) and **D.** composite scores of total BBB permeability in male and female spontaneously hypertensive rats compared to Wistar rats. Chronic hypertension had no effect on permeability in males, however, permeability to all tracers and total BBB score was significantly increased in female SHR compared to normotensive Wistar rats. *p<0.05, **p<0.01; n=7-8/group. Data are mean ± SEM and analyzed by two-way ANOVA with a post-hoc Tukey test to correct for multiple comparisons.

During an open field arena task, there were main effects of sex and hypertension, and a significant interaction between sex and hypertension on locomotion, including speed (F_1,28_ = 19.14; p=0.0002; F_1,28_ = 9.346; p=0.0051; F_1,28_ = 7.388; p=0.0115 by two-way ANOVA, respectively) and distance traveled (F_1,28_ = 18.92; p=0.0002; F_1,28_ = 9.280; p=0.0053; F_1,28_ = 7.112; p=0.0130 by two-way ANOVA, respectively). Compared to all other groups, female Wistar rats had higher average speed (male Wistar: 0.042 ± 0.004 m/s, male SHR: 0.041 ± 0.002 m/s, female Wistar: 0.065 ± 0.005 m/s, female SHR: 0.046 ± 0.001 m/s by two-way ANOVA with post-hoc Tukey’s test) and traveled a greater distance (male Wistar: 12.50 ± 1.36 m, male SHR: 12.12 ± 0.57 m, female Wistar: 19.51 ± 1.48 m, female SHR: 13.80 ± 0.43 m by two-way ANOVA with post-hoc Tukey’s test).

The ventral hippocampus is involved in mediating anxiety-like behavior and is involved in neuropsychiatric disorders, including anxiety. To understand how chronic hypertension impacted anxiety levels, and whether this differed by sex, we assessed several anxiety-like behaviors during an open field test, including thigmotaxis (preferentially remaining near a solid wall) and risk assessment behavior. In rodents, heightened anxiety is demonstrated by a longer latency and fewer entrances/crosses through the center of an arena, and vice versa. There was a main effect of hypertension, but not sex, and no significant interaction between the two on latency to cross the center of the arena (F_1,28_ = 4.832; p=0.0364; F_1,28_ = 2.239; p=0.1458; F_1,28_ = 0.3023; p=0.5868 by two-way ANOVA, respectively). Male and female SHR had decreased latency to the center of the arena compared to normotensive rats, however, this did not reach statistical significance in post-hoc analyses (p>0.05 by two-way ANOVA with post-hoc Tukey test; Figure 2A). There were main effects of sex and hypertension, and no significant interaction between the two on the number of crossings through the center of the arena (F_1,28_ = 9.099; p=0.0054; F_1,28_ = 8.281; p=0.0076; F_1,28_ = 0.2032; p=0.6557 by two-way ANOVA, respectively), although no significant differences were revealed upon completion of post-hoc analyses (Figure 2B).

**Figure 2.**
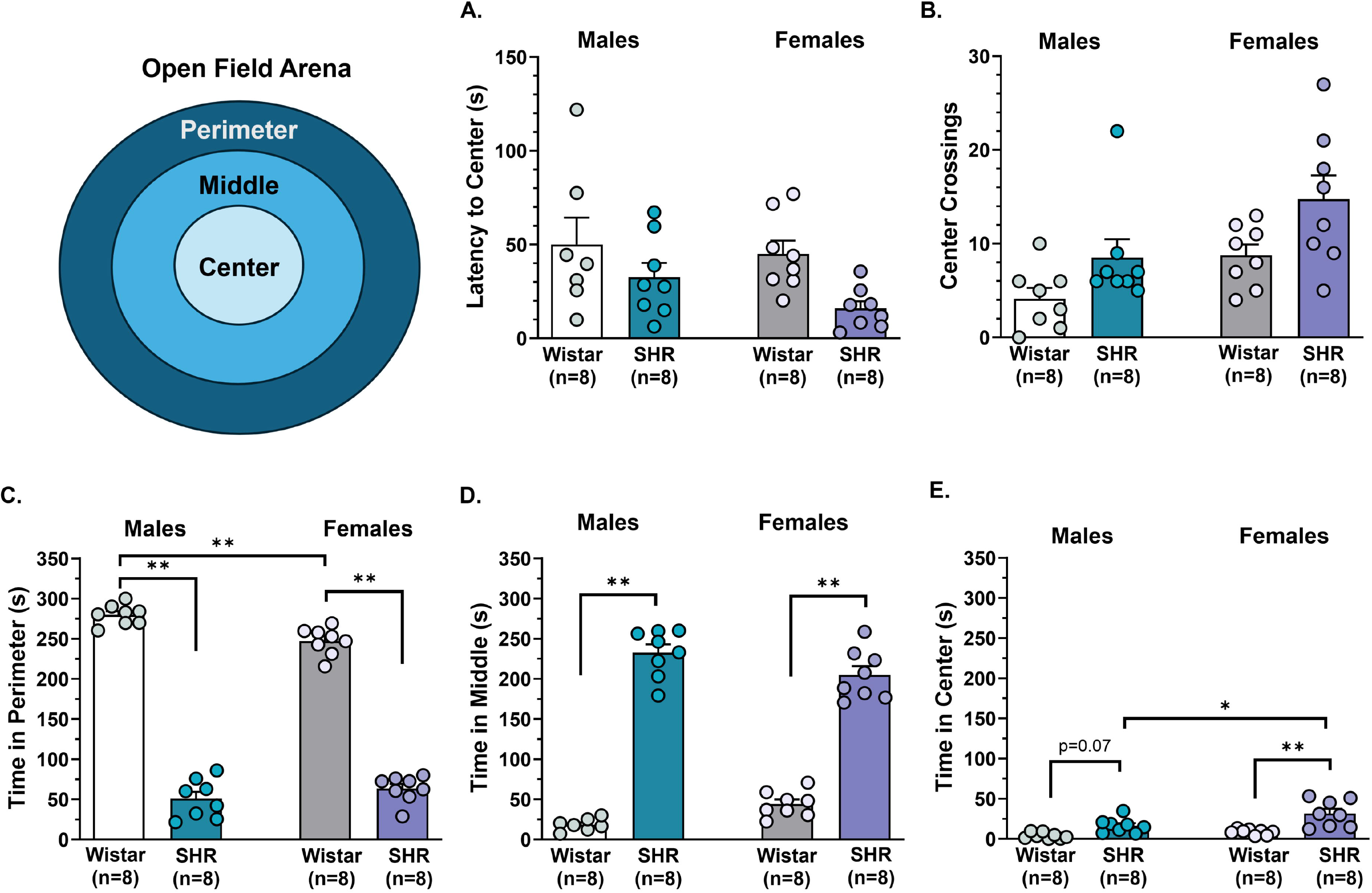
Exploratory behavior and thigmotaxis of male and female spontaneously hypertensive rats (SHR) and normotensive Wistar rats. **A.** Latency to cross through the center of the open field arena and **B.** the number of center crossings in normotensive and hypertensive rats of both sexes. The amount of time spent **C.** in the perimeter, **D.** the middle and **E.** the center of the arena as an assessment of thigmotaxis during the 300 second (5 minute) open field task. *p<0.05, **p<0.01; n=8/group. Data are mean ± SEM and analyzed by two-way ANOVA with a post-hoc Tukey test to correct for multiple comparisons.

To better understand anxiety-like behavior as it relates to thigmotaxis, we quantified the amount of time each rat spent in the outer-most (perimeter; Figure 2C), middle (Figure 2D), and inner-most (center; Figure 2E) of the arena during a 5 min (300 sec) open field test. Increased anxiety-like behavior is associated with more time in the perimeter of the arena, whereas lower anxiety-like behavior consists of more time spent in the middle and inner portions of the arena. There was a main effect of hypertension, but not sex, and a significant interaction between hypertension and sex on the amount of time spent in the perimeter (F_1,28_ = 1048; p<0.001; F_1,28_ = 2.505; p=0.1247; F_1,28_ = 12.58; p=0.0014 by two-way ANOVA, respectively). Normotensive male and female rats spent much of the time in the outer portion of the arena, although females spent less time than males (p<0.01 by two-way ANOVA with post-hoc Tukey test; Figure 2C). SHR, however, spent substantially less time in the perimeter of the arena compared to Wistar rats (p<0.01 by two-way ANOVA with post-hoc Tukey test; Figure 2C). In contrast, SHR of both sexes spent much of the time in the middle zone of the arena, as there were main effects of hypertension, but not sex, and a significant interaction between the two on time in the middle (F_1,27_ = 491.9; p<0.001; F_1,27_ = 0.0014; p=0.9078; F_1,27_ = 9.909; p=0.0040 by two-way ANOVA, respectively). There were main effects of both sex and hypertension on the time spent in the center of the arena, but no significant interaction between the two (F_1,28_ = 8.266; p=0.0076; F_1,28_ = 26.35; p<0.0001; F_1,28_ = 2.261; p=0.1439 by two-way ANOVA, respectively). Female SHR spent more time in the center of the arena compared to female Wistar rats and male SHR (p<0.05 by two-way ANOVA with post-hoc Tukey test; Figure 2E). Male SHR trended towards spending more time in the center of the arena compared to male Wistar rats (p=0.07 by two-way ANOVA with post-hoc Tukey test; Figure 2E). Together, these findings suggest that chronic hypertension had a general anxiolytic effect on thigmotaxis, however, this occurred to a greater extent in females compared to males.

Rearing is a natural component of exploring a novel environment for rodents, and increased levels of anxiety are known to decrease both supported rearing (i.e., using a wall) and unsupported rearing (out in the open). There were main effects of sex, hypertension, and a significant interaction between sex and hypertension on the amount of supported rearing during an open field test (F_1,28_ = 5.947; p=0.0213; F_1,28_ = 7.918; p=0.0089; F_1,28_ = 17.34; p=0.0003 by two-way ANOVA, respectively). Normotensive female rats explored via supported rearing more times than normotensive males and female SHR (p<0.01 by two-way ANOVA with post-hoc Tukey test; Figure 3A); however, there were no main effects or an interaction between sex and hypertension on the amount of time spent performing supported rearing (p>0.05; Figure 3B). There was a main effect of chronic hypertension but not of sex, and no significant interaction between the two on the number of unsupported rears (F_1,28_ = 19.75; p=0.0001; F_1,28_ = 0.2955; p=0.5910; F_1,28_ = 0.0828; p=0.7756; Figure 3C) and amount of time spent rearing during an open field test (F_1,28_ = 23.53; p<0.0001; F_1,28_ = 2.064; p=0.1619; F_1,28_ = 0.6418; p=0.4298 by two-way ANOVA, respectively; Figure 3D). SHR of both sexes reared more and spent more time rearing in the arena without the support of the wall compared to normotensive rats (p<0.05 by two-way ANOVA with post-hoc Tukey test). These findings suggest that despite more supported rearing in normotensive Wistar rats, there was no effect of sex or hypertension on supported rearing, as it contributes to general exploratory behavior. However, there was increased unsupported rearing in both male and female SHR, suggesting an anxiolytic effect of chronic hypertension and increased active exploration.

**Figure 3.**
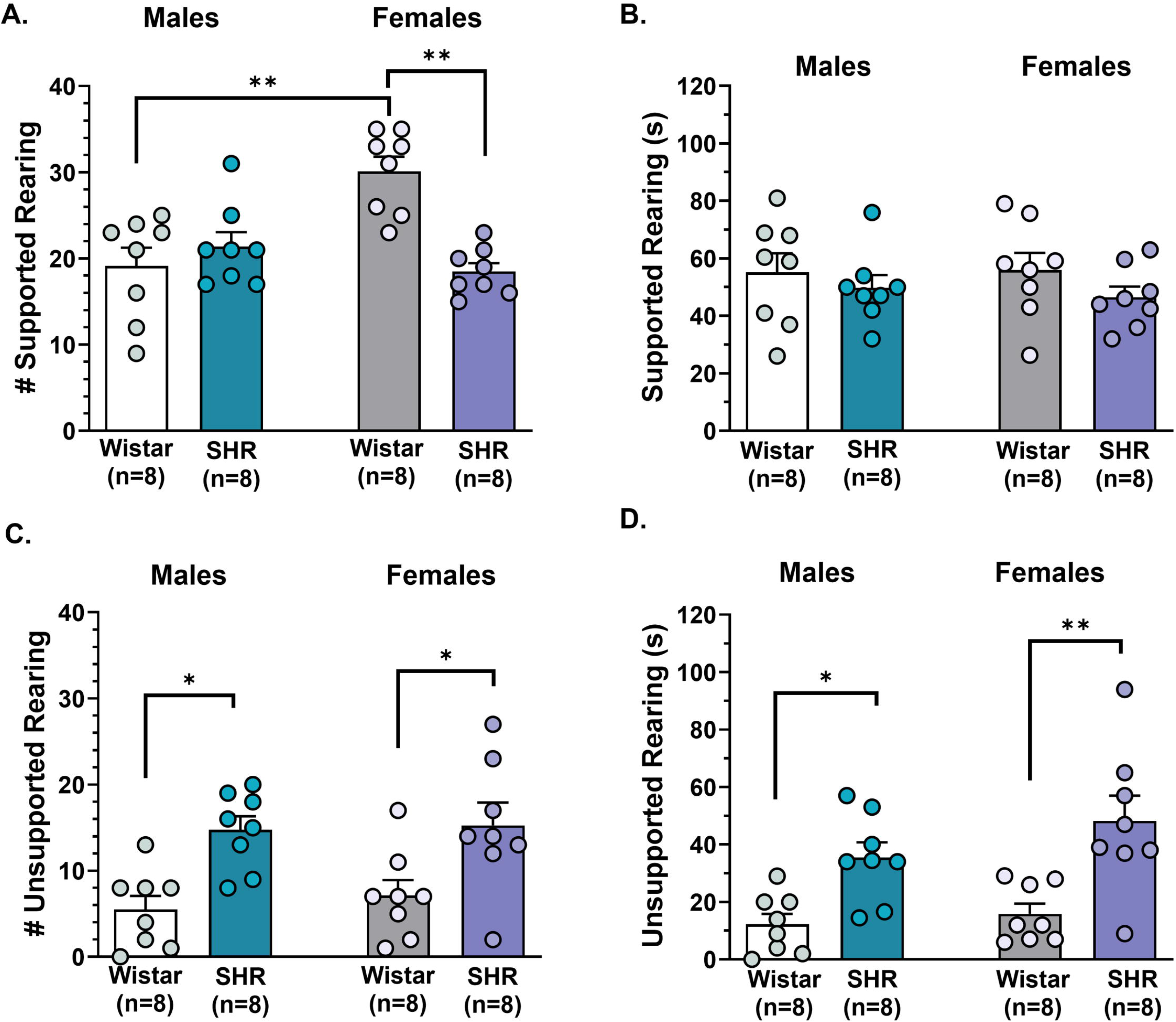
Rearing behavior of male and female spontaneously hypertensive rats (SHR) and normotensive Wistar rats. **A.** The number of supported rears and **B.** time spent rearing with wall support, and **C.** number of unsupported rearing and **D.** time spent rearing in the arena without wall support during the open field task. *p<0.05, **p<0.01; n=8/group. Data are mean ± SEM and analyzed by two-way ANOVA with a post-hoc Tukey test to correct for multiple comparisons.

Risk assessment is closely associated with exploratory behavior and influenced by the level of anxiety in rodents when in a novel environment. Therefore, we quantified the number and amount of time rats demonstrated stretch-attend posture, stretching from the perimeter of the arena towards the center of the arena. Figure 4 shows that there was a main effect of hypertension, but not sex, and a significant interaction between hypertension and sex on amount of stretch-attend posture, as indicated by the number of stretches toward the center of the arena (F_1,28_ = 10.97; p=0.0026; F_1,28_ = 3.387; p=0.0763; F_1,28_ = 4.351; p=0.0462 by two-way ANOVA, respectively; Figure 4A). Further, there were main effects of hypertension and sex, and a significant interaction between the two on the time spent in stretch-attend posture (F_1,28_ = 11.27; p=0.0023; F_1,28_ = 5.875; p=0.0221; F_1,28_ = 5.875; p=0.0221 by two-way ANOVA, respectively; Figure 4B). Female SHR demonstrated significantly more stretch-attend posturing and spent more time stretching to the center compared to normotensive females and hypertensive males (p<0.05 by two-way ANOVA with post-hoc Tukey test; Figure 4A and 4B). These data suggest that there was more approach-avoidance conflict and risk assessment behavior during chronic hypertension in females, but not males.

**Figure 4.**
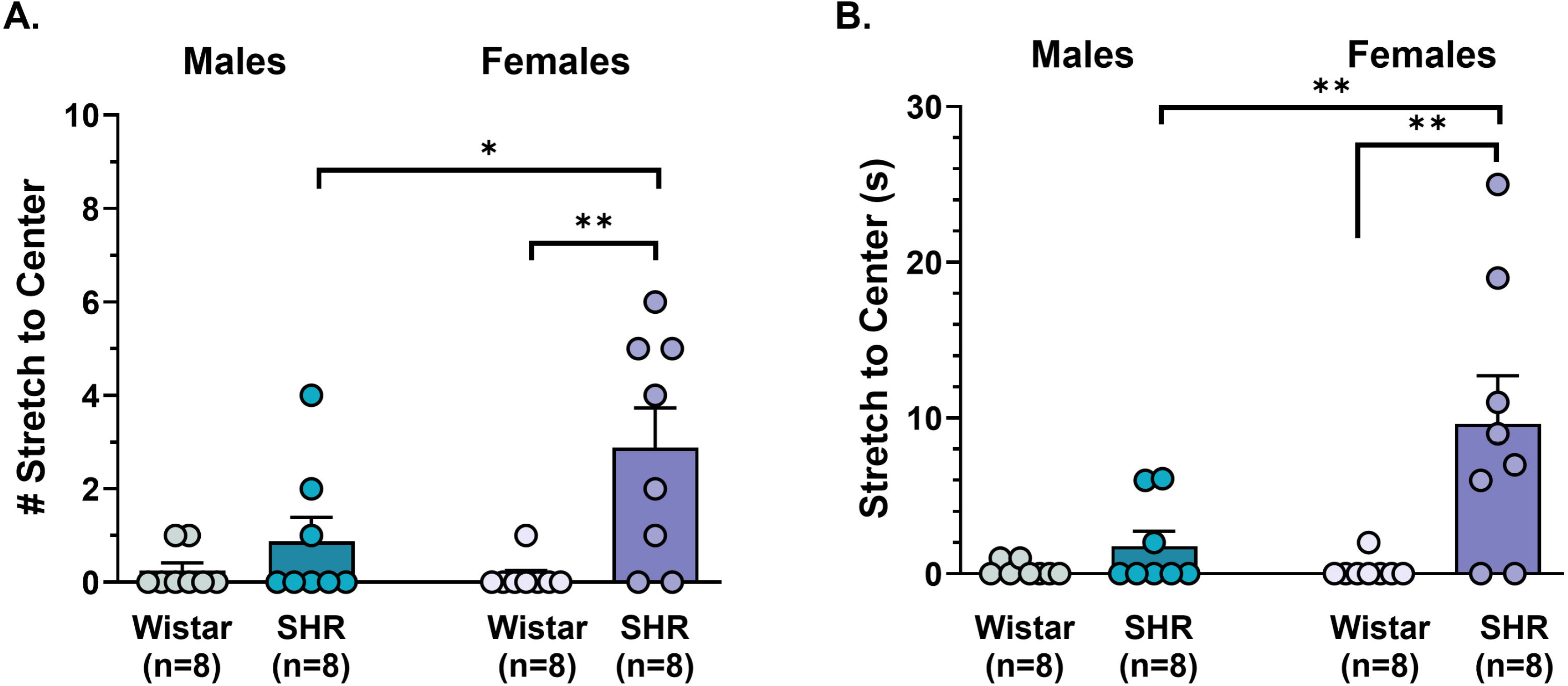
Stretch-attend posture of risk assessment in an open field arena. **A.** The number of times male and female normotensive Wistar rats and spontaneously hypertensive rats (SHR) stretched towards the center of the arena and **B.** the total time spent in this posture during an open field task. *p<0.05, **p<0.01; n=8/group. Data are mean ± SEM and analyzed by two-way ANOVA with a post-hoc Tukey test to correct for multiple comparisons.

Anxiety-like behaviors are controlled by specific ventral hippocampal circuits that may be differentially disrupted by increased BBB permeability during chronic hypertension. To understand the relationship between BBB permeability and anxiety-like behaviors correlation matrices were generated for each sized fluorescent tracer and total BBB permeability score in male and female rats (Figure 5). In males, in general, more permeability was associated with increased anxiety-like behavior, including longer duration to cross the center and crossing fewer times, although there were no statistically significant correlations (Figure 5A). In contrast, increased BBB permeability in the ventral hippocampus of females was negatively correlated with latency to the center and positively correlated with a greater number of crossings through the center, indicating permeability had an anxiolytic effect during hypertension in females. Interestingly, increased BBB permeability was also positively correlated with stretch-attend behavior (Figure 5B), suggesting SHR had higher risk assessment than normotensive females and that BBB permeability affected anxiety differently based upon the specific form of anxiety-like behavior.

**Figure 5.**
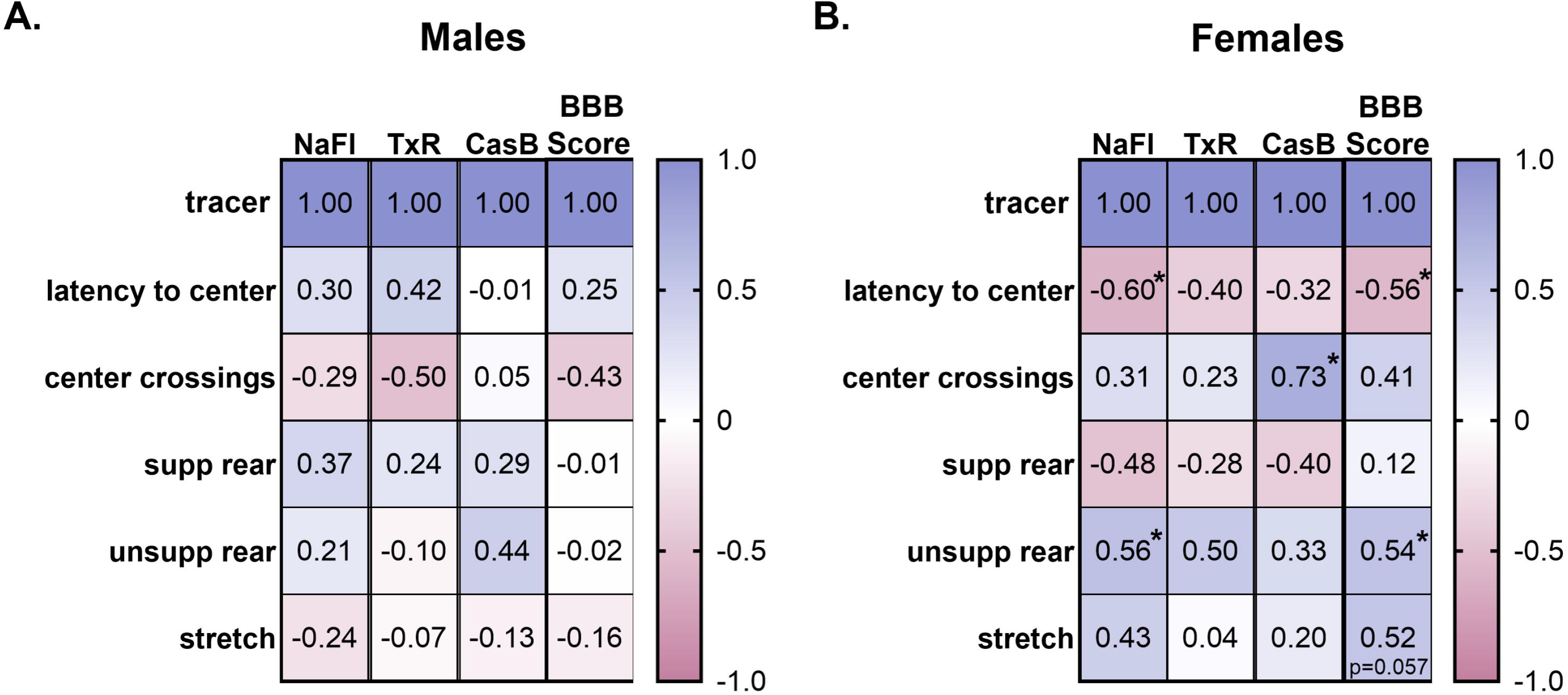
Relationship between blood-brain barrier (BBB) permeability and anxiety-like behavior. Multi-linear regression analyses were performed to understand the relationship between BBB permeability in ventral hippocampus to sodium fluorescein (NaFl), 3 kDa Texas red (TxR), 10 kDa cascade blue (CasB) and total BBB permeability score and exploratory and anxiety-like behavior of **A.** male and **B.** female normotensive Wistar rats and spontaneously hypertensive rats (SHR). Data are mean ± SEM. *p<0.05 by multi-linear regression analyses with Pearson’s correlations.

The BBB is a dynamic interface between what is circulating systemically and the delicate brain milieu. To understand potential mechanisms by which chronic hypertension damages the BBB in females but not males, serum factors known to increase BBB permeability were measured in male and female SHR using ELISAs. Circulating VEGF and IL-17A were elevated in serum from female SHR, although there were no differences in serum ICAM-1 levels and MMP-9 was higher in males than females (Table 1; p<0.01 by Welch’s t-test). In addition, circulating GFAP and NFL were higher in female SHR compared to males, indicating greater neuronal and glial injury as well as increased BBB disruption during chronic hypertension in the female sex (Table 1).

**Table 1.** Circulating factors during chronic hypertension.

|  | **Male SHR (n=6)** | **Female SHR (n=6)** |
|---|---|---|
| VEGF (pg/mL) | 18.91 ± 4.02 | 39.23 ± 4.00** |
| ICAM-1 (ng/mL) | 26.98 ± 2.94 | 24.23 ± 1.28 |
| IL-17A (pg/mL) | 20.31 ± 0.50 | 22.36 ± 0.15** |
| MMP-9 (pg/mL) | 8.76 ± 0.59** | 5.81 ± 0.48 |
| GFAP (pg/mL) | 0.90 ± 0.31 | 1.81 ± 0.25* |
| NFL (pg/mL) | 7.55 ± 1.74 | 28.3 ± 5.31** |

Data are mean ± SEM. \*p<0.05, \*\*p<0.01 by Welch’s t-test. n = 6/group.

## Discussion

Chronic hypertension is an established risk factor for cerebrovascular damage in the neocortex and accelerates age-related cognitive decline.^1^ However, the extent to which hypertension impacts the cerebrovasculature in subcortical limbic brain regions and the relationship between hypertension and neuropsychiatric disorders, remains unclear. In the current study, we compared BBB permeability in the ventral hippocampus in both males and females. While there were no baseline sex differences under healthy conditions, we found that chronic hypertension increased circulating levels of permeability factors and caused BBB disruption in a sex-dependent manner. Females, but not males, had substantially increased extravasation of dextran into the ventral hippocampus during hypertension compared to normotensive females and hypertensive males. This female-specific BBB dysfunction was differentially correlated with anxiety-like behavior, reducing exploration-related anxiety (e.g., thigmotaxis) but augmenting risk assessment behavior that may be due to neuronal injury and/or neuroinflammation. Interestingly, males demonstrated few changes in anxiety-like behavior during chronic hypertension. Overall, this study advances our understanding of how chronic hypertension disrupts the cerebral circulation to include deep limbic brain regions. Further, these findings identify the ventral hippocampal BBB as a site of vulnerability to hypertension-related injury that could contribute to neuropsychiatric disorders in the female sex.

Anxiety can cause transient elevations in blood pressure (i.e., “white coat” hypertension), although it does not, per se, cause chronic hypertension. However, chronic hypertension and anxiety often co-exist, making understanding a potential relationship important when considering mental and cardiovascular well-being. In contrast to our hypothesis, the current study shows that BBB permeability was increased in the ventral hippocampus during chronic hypertension exclusively in the female sex that may be due to higher circulating levels of factors known to damage the BBB and increase paracellular permeability. IL-17A is a proinflammatory cytokine involved in regulating immune responses that promotes neuroinflammation and sympathetic excitation in autonomic brain regions and causes hypertension.^24–26^ It gains access to the brain by causing disruption of the BBB, specifically via degradation of tight junction protein occludin, and circulating levels of IL-17A positively correlated with extravasation of a 10 kDa FITC into the paraventricular nucleus of the hypothalamus.^26^ Thus, higher serum levels of IL-17A in female compared to male SHR could contribute to increased BBB permeability in the ventral hippocampus. Whether levels of IL-17A were elevated above normotensive levels in males that would suggest either a biologically relevant threshold for BBB disruption or increased sensitivity of the BBB to IL-17A-mediated disruption in females remains unclear.

In the current study, males had higher circulating MMP-9 and similar levels of ICAM-1 as in females, both of which are known to degrade tight junction proteins and increase BBB permeability.^27, 28^ Thus, the BBB in the ventral hippocampus may be more sensitive to disruption during chronic hypertension in the female sex, and/or there may be region-specific damage at the BBB during chronic hypertension. In fact, a recent study of ours investigating how social isolation in adulthood compounded chronic hypertension-induced BBB disruption in cognition-centric brain regions showed BBB disruption occurred predominantly in dorsal hippocampus in males and medial prefrontal cortex (mPFC) in females.^18^ That there was no increase in BBB permeability in the ventral hippocampus of males in the current study suggests the effects of chronic hypertension on BBB integrity differs along the septotemporal axis of the hippocampal formation, with disruption occurring in the dorsal hippocampus in hypertensive males and in the ventral hippocampus in hypertensive females. Alternatively, if there are sex-specific microvascular density differences such that females have more robust vascularization of the ventral hippocampus than males during hypertension, such increased surface area could contribute to increased permeability in the female sex. Regardless, these regional differences in hippocampal BBB disruption may contribute to the propensity of anxiety and other ventral-hippocampal-mediated neuropsychiatric disorders to occur in the female sex.

In addition to proteins such as tight junctions forming a structural barrier between circulating factors and the brain, there is also a critical biochemical/selective barrier comprised of specific transport proteins known as efflux transporters.^29^ BBB efflux transporters are powerful ATP-driven pumps that prevent passage of ligands (i.e., cytokines, xenobiotics, etc.) into the brain by actively effluxing them back into the circulation.^30–32^ In fact, NaFl, the smallest tracer used in the current study to measure BBB permeability *in vivo*, is a ligand of BBB efflux transporters.^33^ Therefore, hypertension-related decreases in efflux transporter expression and/or activity at the BBB may contribute to the increased passage of tracers into the ventral hippocampus in the current study. While cytokines elevated to a greater extent in female than male SHR (e.g., IL-17A) are known to increase BBB permeability via degradation of tight junction proteins, additional studies are needed to understand if functional aspects of the BBB are compromised in a sex-specific manner during chronic hypertension.^29, 32^ Further, cell type-specific contributions to BBB damage occurring during chronic hypertension in females remain to be determined (e.g., pericytes, astrocytic endfeet).

The ventral hippocampus is centrally involved in limbic circuits and known to bidirectionally influence anxiety metrics, both increasing anxiety and abrogating it.^17^ The presence and absence of activity in the ventral hippocampus represents high or low anxiety states. Here, we demonstrated selective BBB disruption in the female ventral hippocampus that correlated with anxiety-like behavior with little-to-no relationship between BBB permeability and anxiety-like behavior in males. This is in contrast to lesion studies that have revealed a larger role of the ventral hippocampus in control of anxiety-like behavior in otherwise healthy males compared to females.^34^ Regardless, in conjunction with our previous study showing increased BBB permeability in medial prefrontal cortex in female SHR, these findings suggests that ventral hippocampal – mPFC neurocircuitry is selectively disrupted during chronic hypertension. This could disrupt emotional behavior and cause dysregulation of anxiety and aversion, fear conditioning and extinction and impaired social memory and behavior.^17^

SHR are a commonly used model of attention deficit/hyperactivity disorder during early life (e.g., adolescence) with high levels of locomotion and impaired working memory.^16,35^ In adulthood, persistently increased locomotion can reduce several anxiety-related parameters, such as latency to cross the center of an open field arena. Previous studies in adult SHR have reported general anxiolytic effects using open field tasks and elevated plus maze tasks, although the majority were conducted only in males.^15, 36, 37^ One study that included both sexes reported anxiolytic effects to a similar degree in adult male and female SHR as assessed using an elevated plus maze.^38^ Results from the current study showing significantly reduced thigmotaxis in SHR compared to normotensive rats supports these prior studies. Here, we also show that risk assessment behavior, assessed via stretch-attend posture, was increased in female but not male SHR.^20^ Importantly, locomotion of male and female SHR was similar to normotensive males and lower than normotensive females, suggesting hypertension-related changes in behavior were not due to locomotion differences but more likely due to changes in anxiety.

This study investigated the effects of chronic hypertension and sex on BBB integrity in the ventral hippocampus and its relationship with anxiety-related behavior. However, there are several important limitations of the current study. For example, we used an open field task to assess exploration, thigmotaxis, and risk assessment and found that chronic hypertension differentially affected these behaviors, with more pronounced effects in females. Additional assessments utilizing more anxiety-specific tasks would provide a more thorough investigation of anxiety-related behaviors (i.e., elevated plus maze, successive alleys test). We used a rat model of chronic hypertension and vascular cognitive impairment to understand the relationship between cerebrovascular disease and emotional processing and whether this differed by sex. Studies often investigate potential compounding effects of an additional stressor such as social isolation or chronic unpredictable stress, which could exacerbate BBB damage and anxiety-like behavior during chronic hypertension. In fact, although baseline anxiety levels have been reported to be lower in adult SHR, studies suggest that severe chronic stressors (e.g., chronic restraint stress) cause exaggerated anxiety- and depressive-like responses. The current study did not include molecular assays to measure mechanisms linking BBB damage to behavior, such as neuroinflammation or neuronal network (dys)function. This was due to the necessity of the use of the brain tissue in quantification of extravasation of tracers through the BBB. However, that female SHR had higher circulating levels of GFAP and NF-L supports increased BBB disruption allowing such proteins constrained to the central nervous system under healthy conditions. In addition, this demonstrates increased neurodegeneration and astrocytic activation during hypertension in females compared to males.^39^ As these was measured systemically in serum, future studies are needed to determine local neuroglial injury in the ventral hippocampus during hypertension and its relationship (e.g., cause vs. consequence) with BBB permeability and emotional behavior.

## Conclusions

Overall, we report that BBB permeability in the ventral hippocampus was increased during chronic hypertension in females, but not in males. This BBB disruption was associated with higher levels of circulating permeability factors and biomarkers of neurodegeneration and correlated with anxiety-like behavior. This study advances our understanding of underlying mechanisms by which the female sex may be more susceptible to neuropsychiatric disorders involving ventral hippocampal function in the setting of cardiovascular diseases. The sex-specific increase in BBB permeability may be particularly important in later life once females become post-menopausal and protective effects of estrogens on cerebral endothelial cells, the BBB, and neurons and glia are lost.^40, 41^

## Abbreviations

AAALAC: Association for Assessment and Accreditation of Laboratory Animal Care International
ANOVA: analysis of variance
BBB: blood-brain barrier
CasB: cascade blue
ELISA: enzyme-linked immunosorbent assay
GFAP: glial fibrillary acidic protein
IACUC: Institutional Animal Care and Use Committee
ICAM-1: intercellular adhesion molecule-1
IL-17A: interleukin-17A
NaFl: sodium fluorescein
NF-L: neurofilament light chain
NIH: National Institutes of Health
MMP-9: matrix metalloproteinase-9
mPFC: medial prefrontal cortex
PBS: phosphate buffered saline
RFU: relative fluorescent unit
SEM: standard error of the mean
SHR: spontaneously hypertensive rats
TCA: trichloroacetic acid
TxR: Texas red dextran
VEGF: vascular endothelial growth factor

## Declarations

### Ethics approval and consent to participate

All procedures were approved by the Binghamton University – State University of New York Institutional Animal Care and Use Committee (IACUC Protocol #s 898-23 and 956-25; PI: Chapman) and conducted in accordance with the NIH Guide for the Care and Use of Laboratory Animals and according to the ARRIVE 2.0 guidelines

## Consent for publication

Not applicable.

## Availability of data and materials

All datasets generated and/or analyzed during the current study are publicly available within this manuscript and in the associated Harvard Dataverse repository (https://doi.org/10.7910/DVN/SIFCDV).

## Authors’ contributions

JL was involved in the study design, constructing figures, and writing the initial draft of the manuscript. GMD was involved in conducting permeability studies, constructing figures, statistical analyses and writing the initial draft of the manuscript. AT was involved in conducting permeability studies, performing ELISAs, constructing figures, statistical analyses and funding acquisition. GAB, SEC, EDS and DC were involved in behavioral assessments and analyses. ACC was involved in conceptualization, study design, funding acquisition, supervision, resources, statistical analyses, constructing figures, and writing the initial draft of the manuscript. All Authors read and approved the final manuscript.

## Acknowledgements

Not applicable.

## References

1. Pacholko A and Iadecola C. Hypertension, Neurodegeneration, and Cognitive Decline. Hypertension 2024; 81: 991–1007. 20240301. DOI: 10.1161/HYPERTENSIONAHA.123.21356.

2. Sorond FA, Gorelick PB, Bae HJ, et al. Vascular Contributions to Cognitive Impairment and Brain Health: Clinical Update and Mechanistic Considerations: A Scientific Statement From the American Heart Association. Stroke 2026 20260526. DOI: 10.1161/STR.0000000000000524.

3. Whelton PK, Carey RM, Aronow WS, et al. 2017 ACC/AHA/AAPA/ABC/ACPM/AGS/APhA/ASH/ASPC/NMA/PCNA Guideline for the Prevention, Detection, Evaluation, and Management of High Blood Pressure in Adults: A Report of the American College of Cardiology/American Heart Association Task Force on Clinical Practice Guidelines. Hypertension 2018; 71: e13–e115. 2017/11/15. DOI: 10.1161/hyp.0000000000000065.

4. Cotrell V, Wild K and Bader T. Medication management and adherence among cognitively impaired older adults. J Gerontol Soc Work 2006; 47: 31–46. DOI: 10.1300/J083v47n03_03.

5. Smith D, Lovell J, Weller C, et al. A systematic review of medication non-adherence in persons with dementia or cognitive impairment. PloS one 2017; 12: e0170651. 20170206. DOI: 10.1371/journal.pone.0170651.

6. Walker KA, Sharrett AR, Wu A, et al. Association of Midlife to Late-Life Blood Pressure Patterns With Incident Dementia. Jama 2019; 322: 535–545. DOI: 10.1001/jama.2019.10575.

7. Updated trends in the global prevalence and burden of mental disorders, 1990-2023: a systematic analysis for the Global Burden of Disease Study 2023. *Lancet* 2026; 407: 2040-2064. DOI: 10.1016/s0140-6736(26)00519-2.

8. Zlokovic BV. The blood-brain barrier in health and chronic neurodegenerative disorders. Neuron 2008; 57: 178–201. 2008/01/25. DOI: S0896-6273(08)00034-2 [pii] 10.1016/j.neuron.2008.01.003.

9. Ueno M. Molecular anatomy of the brain endothelial barrier: an overview of the distributional features. Curr Med Chem 2007; 14: 1199–1206. 2007/05/17.

10. Santisteban MM, Iadecola C and Carnevale D. Hypertension, Neurovascular Dysfunction, and Cognitive Impairment. Hypertension 2023; 80: 22–34. 20220921. DOI: 10.1161/HYPERTENSIONAHA.122.18085.

11. Nation DA, Sweeney MD, Montagne A, et al. Blood-brain barrier breakdown is an early biomarker of human cognitive dysfunction. Nature medicine 2019; 25: 270–276. DOI: 10.1038/s41591-018-0297-y.

12. Dion-Albert L, Bandeira Binder L, Daigle B, et al. Sex differences in the blood-brain barrier: Implications for mental health. Front Neuroendocrinol 2022; 65: 100989. 20220307. DOI: 10.1016/j.yfrne.2022.100989.

13. Solarz A, Majcher-Maslanka I and Chocyk A. Effects of early-life stress and sex on blood-brain barrier permeability and integrity in juvenile and adult rats. Dev Neurobiol 2021; 81: 861–876. 20210824. DOI: 10.1002/dneu.22846.

14. Shao X, Shou Q, Felix K, et al. Age-related decline in blood-brain barrier function is more pronounced in males than females in parietal and temporal regions. Elife 2024; 13 20241104. DOI: 10.7554/eLife.96155.

15. Manolova AO, Nedogreeva OA, Mamedova DI, et al. The Relationship between Anxiety, Depression-Like Behavior, and Arterial Pressure: A Comparative Study in 13-Month-Old Wistar, WKY, and SHR Rats. *Advances in Gerontology* 2025; 15: 91-103. DOI: 10.1134/S2079057025600296.

16. Corona JC. Rodent research of attention-deficit/hyperactivity disorder: insights into widely used animal models. Lab Anim Res 2025; 41: 24. 20250923. DOI: 10.1186/s42826-025-00255-5.

17. Biane JS, Wagner-Carena J and Kheirbek MA. The ventral hippocampus: computations, circuits and functions. Nat Rev Neurosci 2026 20260911. DOI: 10.1038/s41583-026-01078-6.

18. Thaqi A, DeMarco GM, Schellinger ED, et al. Chronic hypertension and social isolation disrupt the blood-brain barrier and memory in a sex-dependent manner. Neurobiol Dis 2026: 107563. 20260804. DOI: 10.1016/j.nbd.2026.107563.

19. Johnson AC, Tremble SM, Chan SL, et al. Magnesium sulfate treatment reverses seizure susceptibility and decreases neuroinflammation in a rat model of severe preeclampsia. PloS one 2014; 9: e113670. 2014/11/20. DOI: 10.1371/journal.pone.0113670PONE-D-14-43811 [pii].

20. Sturman O, Germain PL and Bohacek J. Exploratory rearing: a context- and stress-sensitive behavior recorded in the open-field test. Stress 2018; 21: 443–452. 20180216. DOI: 10.1080/10253890.2018.1438405.

21. Holly KS, Orndorff CO and Murray TA. MATSAP: An automated analysis of stretch-attend posture in rodent behavioral experiments. Sci Rep 2016; 6: 31286. 20160809. DOI: 10.1038/srep31286.

22. Johnson AC, Li Z, Orfila JE, et al. Hippocampal network dysfunction as a mechanism of early-onset dementia after preeclampsia and eclampsia. Progress in neurobiology 2021; 199: 101938. DOI: 10.1016/j.pneurobio.2020.101938.

23. Gannon O, Tremble SM, McGinn C, et al. Angiotensin II-mediated hippocampal hypoperfusion and vascular dysfunction contribute to vascular cognitive impairment in aged hypertensive rats. Alzheimers Dement 2024; 20: 890–903. 2023/10/11. DOI: 10.1002/alz.13491.

24. Robert M and Miossec P. Effects of Interleukin 17 on the cardiovascular system. Autoimmun Rev 2017; 16: 984–991. 20170710. DOI: 10.1016/j.autrev.2017.07.009.

25. Cao Y, Yu Y, Xue B, et al. IL (Interleukin)-17A Acts in the Brain to Drive Neuroinflammation, Sympathetic Activation, and Hypertension. Hypertension 2021; 78: 1450–1462. 20211011. DOI: 10.1161/HYPERTENSIONAHA.121.18219.

26. Yu Y, Weiss RM and Wei SG. Interleukin 17A Contributes to Blood-Brain Barrier Disruption of Hypothalamic Paraventricular Nucleus in Rats With Myocardial Infarction. J Am Heart Assoc 2024; 13: e032533. 20240119. DOI: 10.1161/JAHA.123.032533.

27. Dietrich JB. The adhesion molecule ICAM-1 and its regulation in relation with the blood-brain barrier. J Neuroimmunol 2002; 128: 58–68. DOI: 10.1016/s0165-5728(02)00114-5.

28. Rosenberg GA. Matrix metalloproteinases and their multiple roles in neurodegenerative diseases. Lancet Neurol 2009; 8: 205–216. DOI: 10.1016/S1474-4422(09)70016-X.

29. Miller DS, Bauer B and Hartz AM. Modulation of P-glycoprotein at the blood-brain barrier: opportunities to improve central nervous system pharmacotherapy. Pharmacological reviews 2008; 60: 196–209. 2008/06/19. DOI: pr.107.07109 [pii] 10.1124/pr.107.07109.

30. Begley DJ. ABC transporters and the blood-brain barrier. Curr Pharm Des 2004; 10: 1295–1312. 2004/05/12.

31. Hartz AM, Mahringer A, Miller DS, et al. 17-beta-Estradiol: a powerful modulator of blood-brain barrier BCRP activity. J Cereb Blood Flow Metab 2010; 30: 1742–1755. DOI: 10.1038/jcbfm.2010.36.

32. Hartz AM, Bauer B, Fricker G, et al. Rapid modulation of P-glycoprotein-mediated transport at the blood-brain barrier by tumor necrosis factor-alpha and lipopolysaccharide. Mol Pharmacol 2006; 69: 462–470. 2005/11/10. DOI: mol.105.017954 [pii] 10.1124/mol.105.017954.

33. Schoknecht K and Eilers J. Brain-to-blood transport of fluorescein in vitro. Sci Rep 2024; 14: 25572. 20241026. DOI: 10.1038/s41598-024-77040-2.

34. Wang C, Zhang Y, Shao S, et al. Ventral Hippocampus Modulates Anxiety-Like Behavior in Male But Not Female C57BL/6 J Mice. Neuroscience 2019; 418: 50–58. 20190825. DOI: 10.1016/j.neuroscience.2019.08.032.

35. Kim HB, Kim YJ, Lim HM, et al. Comprehensive behavioral profiling in male spontaneously hypertensive rats: latent trait mapping supports a valid multidomain ADHD model. Transl Psychiatry 2026; 16 20260509. DOI: 10.1038/s41398-026-04099-y.

36. Calzavara MB, Lopez GB, Abilio VC, et al. Role of anxiety levels in memory performance of spontaneously hypertensive rats. Behav Pharmacol 2004; 15: 545–553. DOI: 10.1097/00008877-200412000-00003.

37. Tchekalarova J, Krushovlieva D, Ivanova P, et al. Spontaneously hypertensive rats vs. Wistar Kyoto and Wistar rats: An assessment of anxiety, motor activity, memory performance, and seizure susceptibility. Physiol Behav 2023; 269: 114268. 20230610. DOI: 10.1016/j.physbeh.2023.114268.

38. Ferguson SA and Gray EP. Aging effects on elevated plus maze behavior in spontaneously hypertensive, Wistar-Kyoto and Sprague-Dawley male and female rats. Physiol Behav 2005; 85: 621–628. DOI: 10.1016/j.physbeh.2005.06.009.

39. Pontecorvo MJ, Lu M, Burnham SC, et al. Association of Donanemab Treatment With Exploratory Plasma Biomarkers in Early Symptomatic Alzheimer Disease: A Secondary Analysis of the TRAILBLAZER-ALZ Randomized Clinical Trial. JAMA Neurol 2022; 79: 1250–1259. DOI: 10.1001/jamaneurol.2022.3392.

40. Maggioli E, McArthur S, Mauro C, et al. Estrogen protects the blood-brain barrier from inflammation-induced disruption and increased lymphocyte trafficking. Brain, behavior, and immunity 2016; 51: 212–222. DOI: 10.1016/j.bbi.2015.08.020.

41. Mervosh N and Devi G. Estrogen, menopause, and Alzheimer’s disease: understanding the link to cognitive decline in women. Front Mol Biosci 2025; 12: 1634302. 20250630. DOI: 10.3389/fmolb.2025.1634302.

